# Sensitivity to TDP-43 loss and degradation resistance determine cryptic exon biomarker potential

**DOI:** 10.1101/2025.11.23.689722

**Authors:** Anna-Leigh Brown, Matteo Zanovello, Alla Mikheenko, Dario Dattilo, Flaminia Pellegrini, Sam Bryce-Smith, Francesca Mattedi, Puja R. Mehta, Jose Norberto S. Vargas, NYGC ALS Consortium, Jack Humphrey, Matthew J. Keuss, Pietro Fratta

**Affiliations:** Department of Neuromuscular Diseases, UCL Queen Square Institute of Neurology, UCL, London, UK; UK Dementia Research Institute, UCL, London, UK; Departments of Neuroscience, Genetics and Genomic Sciences, Icahn School of Medicine at Mount Sinai, New York, USA; The Francis Crick Institute, London, UK

**Author notes:** These authors contributed equally to this work (Anna-Leigh Brown, Matteo Zanovello, Alla Mikheenko).

## Abstract

Cryptic splicing caused by TDP-43 proteinopathy is a hallmark of the neurodegenerative diseases amyotrophic lateral sclerosis (ALS) and frontotemporal dementia (FTD). However, which cryptic splicing events (CEs) are the most sensitive to TDP-43 depletion, where CEs localise within cells, and how specific CEs are in human tissues is poorly defined. Analyses of *in vitro* TDP-43 knockdowns and postmortem RNA-seq datasets revealed that a small subset out of thousands of CEs are specific markers for TDP-43 proteinopathy *in vivo*. Nonsense-mediated decay (NMD) masked a portion of CEs, influencing their subcellular localization and detectability in tissue. Dose-dependent TDP-43 depletion identified “early-responsive” CEs, which possess stronger splice sites and denser, more canonical TDP-43 binding motifs. Finally, we developed a composite cryptic burden score that effectively captured TDP-43 pathology across heterogeneous tissues and correlated with regional vulnerability and genetic background. Our work identifies robust biomarkers and offers new insights into TDP-43-mediated splicing dysregulation in neurodegeneration.

## Introduction

TDP-43 (TAR DNA Binding Protein 43 kDa, encoded by *TARDBP*) is a ubiquitously expressed, mainly nuclear DNA- and RNA-binding protein that plays a pivotal role in alternative pre-mRNA splicing^1^ by binding to UG-rich motifs^2^. TDP-43 proteinopathy, defined by cytoplasmic accumulation and nuclear loss, is found in postmortem brain samples in 97% of amyotrophic lateral sclerosis (ALS) cases and around half of frontotemporal dementia (FTD) and Alzheimer’s disease (AD), as well as in the muscles of inclusion body myositis (IBM)^3–6^. TDP-43 nuclear loss prevents it from exerting its function as a splicing repressor causing cryptic intronic sequences to be included in mature transcripts^7^. Cryptic splicing events (CEs) comprise novel cassette exons, 5’ and 3’ exon extensions, exon skipping, intron retention, alternative last exon, and alternative first exon events. CEs can lead to a premature stop codon (PTC) or introduction of a frameshift (FS), ultimately resulting in transcript degradation through nonsense-mediated decay (NMD)^8–12^. CEs can also be in-frame, leading to the incorporation of novel peptide sequences^13,14^. TDP-43 also modulates alternative polyadenylation^15^, and its loss can lead to novel last exons^16,17^ or to the usage of novel polyA sites within introns or the 3’UTR, influencing mRNA stability and protein levels^18–20^. TDP-43 CEs show species and tissue specificity^21–23^, but in light of the preferential involvement of neurons and myocytes in TDP-43 proteinopathies, most of the *in cellulo* models for TDP-43 loss of function have focused on these cell types^21,23^. Different CEs modulated by TDP-43 have emerged as potential therapeutic targets, including *UNC13A*^24^*, STMN2*^25^, and *KCNQ2*^26^. Moreover, the in frame CE in *HDGFL2*, which encodes a novel peptide, is a promising candidate for the development of biomarkers for TDP-43 proteinopathies^13,27,28^.

TDP-43 CEs analysis has relied on RNA-sequencing and bioinformatic tools to quantify the differential splicing usage. By employing these techniques, recent work has tried to characterise general properties of TDP-43 CEs, including their sensitivity to NMD^10^ and their differential expression in tissues affected by TDP-43 proteinopathies^23^. However, a comprehensive analysis of the biological properties of TDP-43 cryptic splicing, covering the impact of NMD on their expression, the cellular localisation of CE-bearing transcripts, the sensitivity of CEs to levels of TDP-43 loss, and the role of sequence features in modulating their inclusion, is lacking.

Here, we leverage extensive bioinformatics analyses on both publicly available and novel *in vitro* TDP-43 knockdown (KD) datasets to define CEs sensitivity to NMD, subcellular localisation, sensitivity to TDP-43 loss, and sequence features. Furthermore, by exploiting post-mortem data from ALS and FTD patients, we identify disease-relevant CEs with potential utility as molecular biomarkers in TDP-43 proteinopathies. Finally, we provide an accompanying interactive web browser to allow researchers to easily explore these CE characteristics, NMD sensitivity, gene expression changes, including nuclear/cytoplasmic distribution, and CE expression patterns in post-mortem tissue.

## Results

### A subset of *in vitro* cryptic splicing is specific to TDP-43 proteinopathy in postmortem tissue

To broadly characterise TDP-43 CE, we assembled a compendium of previously published and newly generated *in vitro* TDP-43 KD datasets, including two from neuroblastoma cell lines, five from iPSC-derived neurons, and a complete knock-out in HeLa cells, and performed both differential expression and splicing analyses (Supplementary Tables 1-2). We set a liberal threshold to identify CEs by retaining all splicing events with a percent spliced in (PSI) in controls < 5% and in the TDP-43 KD >10% in any dataset and identified 3,905 CEs in 2,684 genes (Supplementary Table 2). Despite this large number, the vast majority of CEs were detected in only one dataset (only 650 CEs in 436 genes were found in at least 2 datasets) (Fig. 1A). CEs have been reported to show cell-type specificity ^21,22,29^, but when we analysed CEs called significant in at least six datasets, CEs were detectable in the sequencing data even when our splicing analyses did not classify them as significant (Fig. 1B). In the non-neuronal HeLa dataset, undetected events were largely explained by lack of parent gene expression, except for the cryptic exon in *PXDN* (Fig. 1B).

**Figure 1.**
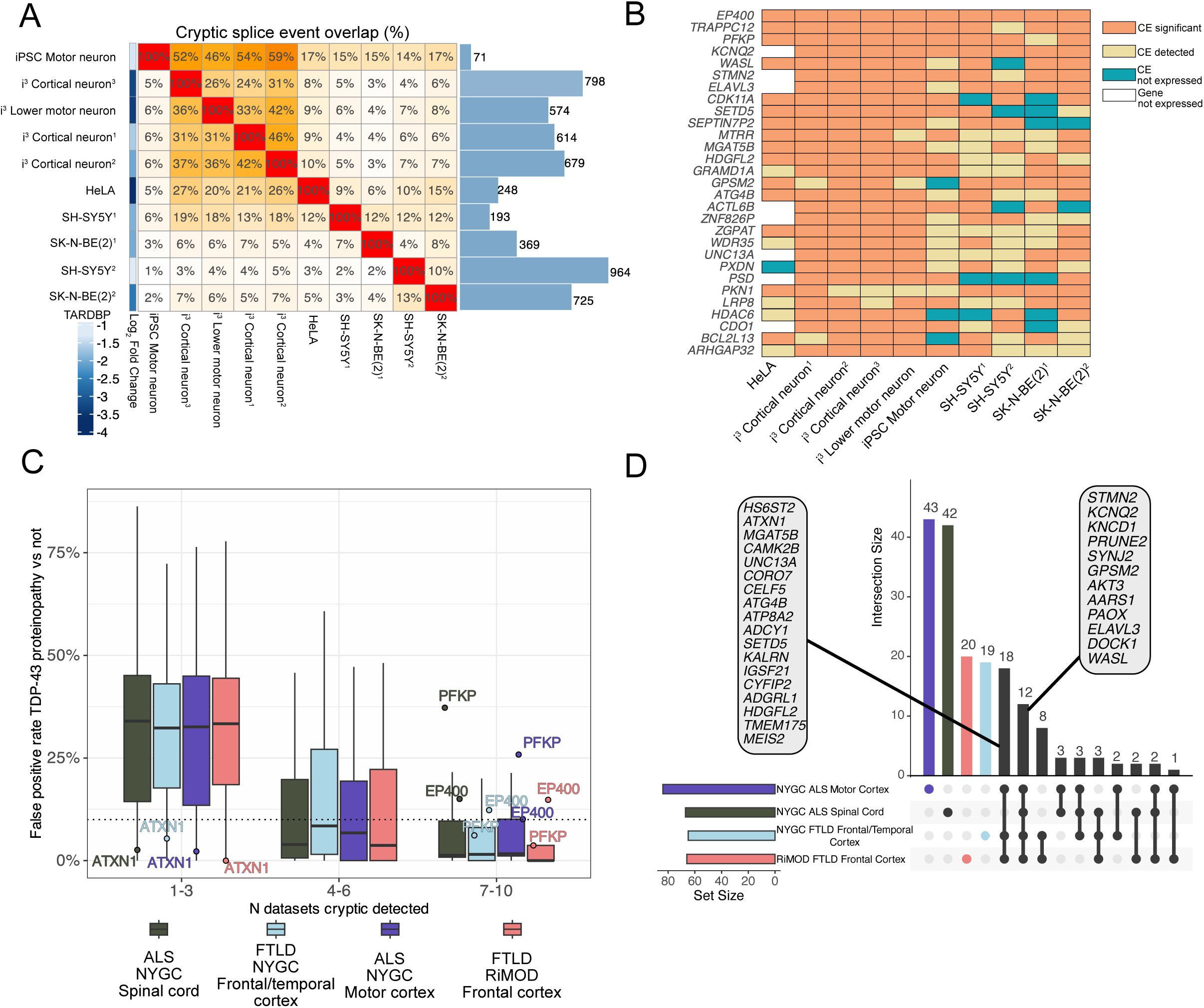
Identification of TDP-43 proteinopathy specific cryptic splicing from in vitro TDP-43 knockdowns. A. Heatmap of overlapping cryptic splice events (CE) across TDP-43 knockdown datasets, clustered by overlap. Bars (right) show the number of CEs (control PSI <5%, knockdown >10%). The left colour scale indicates the log2 fold change of TARDBP expression in knockdown versus control. B. Detection of common CEs across TDP-43 knockdown datasets. CEs present in at least 6 of 10 datasets were manually classified as: (1) significant by splicing software (orange), (2) lowly expressed but not called (>5 junction reads) (yellow), (3) lacking evidence of expression (blue), or (4) absent due to no expression of the parent gene (white). C. False positive rate of CEs in the NYGG and RiMOD datasets based on the number of in vitro datasets where a CE was found. D. Upset plot of genes with a disease-specific CE (FPR < 10%, TPR >10 % and detected in at least five distinct TDP-43 proteinopathy tissue samples). Highlighted genes are those with disease specific CEs present across all datasets and those only in cortical samples.

We next explored if widely expressed CEs were the most promising markers in postmortem tissue for TDP-43 proteinopathy. To this end, we analysed two large postmortem bulk RNA-sequencing datasets including cases with ALS or frontotemporal lobar degeneration (FTLD), the pathological substrate of FTD: the NYGC ALS/FTD consortium (1,945 cortex and spinal cord samples from 457 cases and 121 controls) and the RiMOD-FTD cohort (47 samples from 31 cases and 16 controls). These datasets differ substantially in age at death, RNA integrity, postmortem interval, and library depth (Extended Data Fig. 1A), underscoring the need for CEs that can be reliably detected in heterogeneous post-mortem samples. For each CE detected *in vitro*, we tested whether its z-score–normalised PSI could distinguish control from TDP-43 proteinopathy by reporting the true- and false-positive rates at a decision score of zero, which corresponds to the cohort mean for that event. CEs detected in a single *in vitro* dataset showed poor discriminatory power, typically producing false-positive rates above 10%. A notable exception to this was the *ATXN1* CE, which had an FPR < 2% in any postmortem dataset, despite only being observed in i^3^ lower motor neurons (LMN)s (Fig. 1C). In contrast, recurrent CEs across datasets were stronger predictors, with most showing FPR <10% (Fig. 1C). Notable exceptions to this rule include the *EP400* CE, an exon-skipping event observed in all ten TDP-43 depletion datasets but broadly detected in postmortem tissue not affected by TDP-43 proteinopathy, and the *PFKP* CE, which had a high FPR in motor cortex and spinal cord (Fig. 1C). Detection rate for predictive-CE (FPR ≤ 10%, TPR ≥10 % and detected in at least five distinct TDP-43 proteinopathy tissue samples) showed low correlation with parent gene expression (R = 0.11–0.27; Extended Data Fig. 1B). For example, the *STMN2* CE was the most commonly detected single event in the spinal cord, despite relatively low gene expression in bulk tissue, possibly reflecting *STMN2*’s motor neuron enrichment ^30^.

Even though the majority (3,205 / 3,905) of CEs were detected across any of the postmortem datasets, only 210 met our predictive cutoff for TDP-43 proteinopathy (Supplementary Table 2). We next asked which genes had CEs that met predictive criteria (FPR ≤ 10%, TPR ≥10 % and detected in at least five distinct tissue samples) across all postmortem datasets. Twelve CE-bearing genes met predictive criteria in all postmortem datasets, and eighteen genes had predictive CEs in all three cortical regions. The largest shared overlap occurred between the two FTD cohorts, whereas the ALS spinal cord and motor cortex showed the highest number of unique predictive-CE genes (Fig. 1D). Similar to recently reported work ^23^, we find that baseline gene expression largely did not drive differences in cohort specific biomarker candidates, as predictive-CE genes specific to either the ALS motor cortex or ALS spinal cord were well expressed in either dataset (Extended Data Fig. 1C,D). Finally, we examined predictive-CE expression in neuronal nuclei from ALS–FTD frontal cortices^31^ and found that the majority of predictive-CE from FTD datasets were also upregulated in TDP-43 negative neuronal nuclei, whereas this was the case for a smaller fraction of the predictive-CE found in ALS spinal cord and motor cortex (Extended Data Fig. 1E). Collectively, these data highlight that a small subset of all observed *in vitro* CEs serve as robust RNA-based markers for TDP-43 proteinopathy in postmortem human tissue.

### Nonsense-mediated decay masks dark cryptic splicing

Nonsense mediated decay (NMD) can effectively mask CE-bearing transcripts and inhibition of the NMD machinery can enable their detection, as shown for *UNC13A* CE^8^. To gain insights into the wider impact of NMD on CEs, we performed three orthogonal methods of NMD inhibition treatment with the translational inhibitor cycloheximide (CHX), CRISPRi-mediated KD of the key NMD factor UPF1, and using the small molecule inhibitor of the SMG1 kinase, SMG1-11j. We confirmed NMD inhibition by assessing the expression of known poison exons^32^ (Extended Data Fig. 2A,B). We first classified CEs based on their sensitivity to NMD as either “NMD-rescued” or “non-NMD-rescued.” Across the datasets, events such as *STMN2*, *KCNQ2*, *ACTL6B*, and *HDGFL2* were consistently found to be NMD-insensitive, whereas other cryptic events, such as known NMD-target *ATG4B* and *UNC13A*^7,8^, consistently increased their PSI after NMD-inhibition (Fig. 2A-C). We also identified a subset of “dark cryptic” events detected only under NMD inhibition (Fig. 2A–C; Supplementary Table 2). We found no evidence that TDP-43 KD alone affected NMD directly, as across the public TDP-43 KD datasets we analysed we saw neither upregulation of known poison exons (Extended Data Fig. 2C) nor did we observe UPF1 KD-responsive genes were upregulated following TDP-43 loss (Extended Data Fig. 2D).

**Figure 2.**
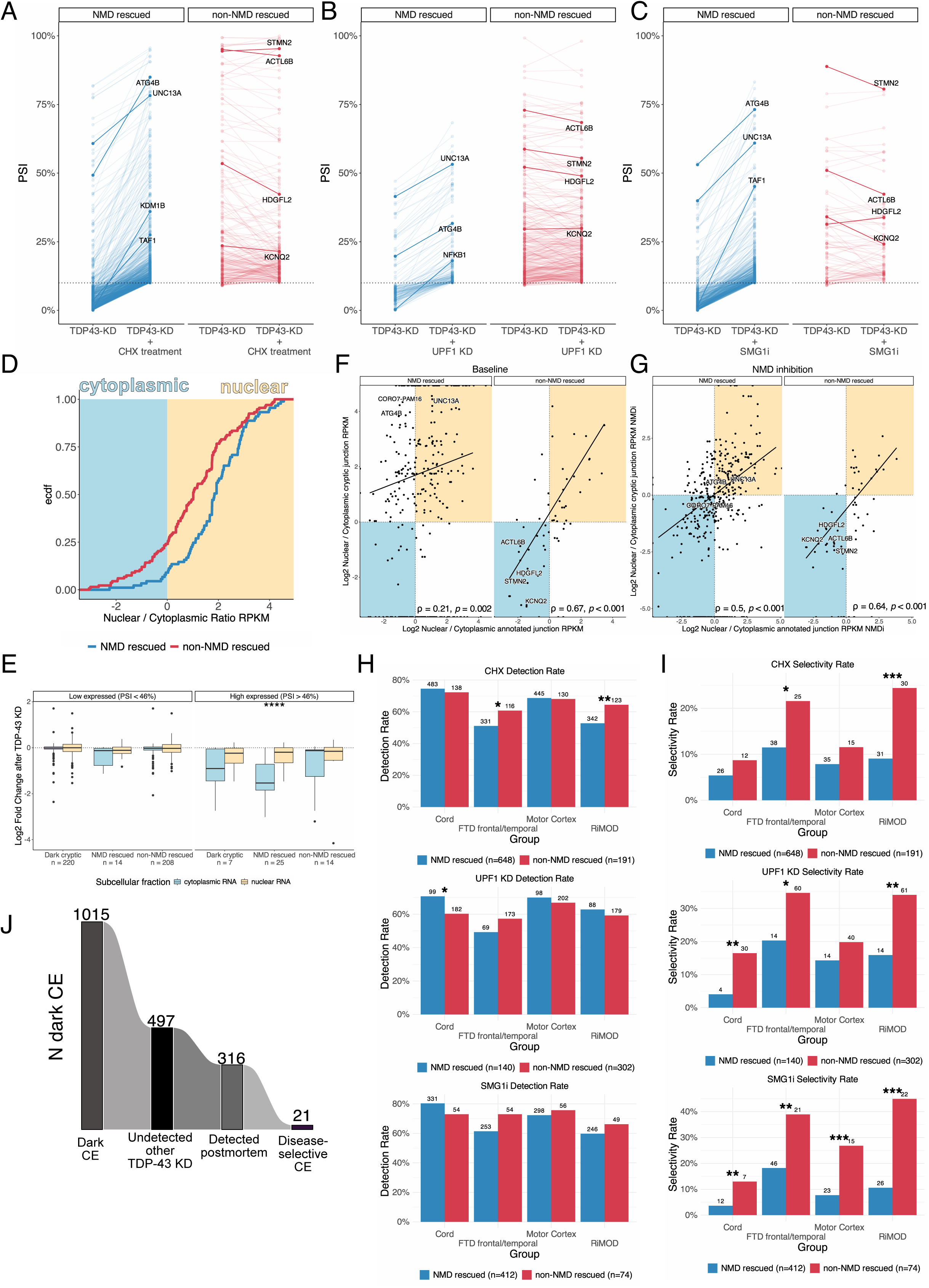
Nonsense-mediated decay affects cryptic splicing detectability, localisation, and selectivity for TDP-43 proteinopathies. A-C) Percent spliced in (PSI) of cryptic splicing events (CE) in (A) SH-SY5Y cells after nonsense-mediated decay (NMD) inhibition by CHX treatment, (B) i3-derived cortical-like neurons after NMD inhibition by UPF1 knockdown, (C) SH-SY5Y cells after NMD inhibition by SMGi-j11, split by genes that are rescued by more than 5% (left, blue) or less (right, red). Highlighted genes include exemplar genes where NMD-sensitivity has been described before and example dark cryptic genes. D. Nuclear to cytoplasmic ratio NMD- (blue) and non-NMD (red) sensitive CEs. E. Gene expression changes after TDP-43 knockdown in genes carrying CEs, divided by NMD category, gene expression, and subcellular fraction. Box plots: boundaries 25–75th percentiles; midline, median; whiskers, Tukey style. Wilcoxon test. F. Scatter plots relating the nuclear to cytoplasmic ratio of annotated junctions (x-axis) and the nuclear to cytoplasmic ratio of cryptic junctions (y-axis) in baseline condition, split between NMD rescued and non-NMD rescued CEs. G. Scatter plots relating the nuclear to cytoplasmic ratio of annotated junctions (x-axis) and the nuclear to cytoplasmic ratio of cryptic junctions (y-axis) after NMD inhibition, split between NMD rescued and non-NMD rescued CEs. H. Detection rate of NMD- (blue) and non-NMD-rescued (red) genes based on the three NMD inhibition experiments in the NYGC ALS motor cortex, NYGC ALS spinal cord, NYGC FTLD frontal/temporal cortex, RiMOD FTLD frontal cortex. Fisher’s exact test. I. Selectivity rate of NMD- (blue) and non-NMD-rescued (red) genes based on the three NMD inhibition experiments in the NYGC ALS motor cortex, NYGC ALS spinal cord, NYGC FTLD frontal/temporal cortex, RiMOD FTLD frontal cortex. Fisher’s exact test. J. Number of dark CEs across three NMD-inhibition datasets, including total number, number undetected in other TDP-43 knockdowns, number observed in postmortem tissue (>2 junction reads), and number disease selective for TDP-43 proteinopathy. Significance levels reported as * (p<0.05) ** (p<0.01) *** (p<0.001) **** (p<0.0001).

### Nonsense-mediated decay impacts cryptic exon localisation and selectivity for TDP-43 proteinopathy

Next we assessed the interplay between NMD and CEs subcellular localisation by performing nuclear and cytoplasmic fractionation under TDP-43 KD with and without NMD inhibition by SMG1-11j treatment. Consistent with NMD-mediated transcript decay primarily occurring in the cytoplasm, NMD-rescued events displayed higher nuclear expression than non-NMD-rescued (Fig. 2D). In line with this cytoplasmic bias, high-PSI, NMD-sensitive cryptic-containing genes exhibited pronounced cytoplasmic depletion (Fig. 2E).

We next compared how the localisation of CE-containing transcripts compared to the localisation of their corresponding annotated isoform. Under baseline conditions, non-NMD-rescued cryptics closely mirrored the nuclear/cytoplasmic distribution of their canonical isoforms (ρ = 0.67, p < 0.001) (Fig. 2F). In contrast, NMD-sensitive CEs generally diverged from their canonical isoforms, with higher nuclear expression than their annotated isoform under baseline conditions. (ρ = 0.21, p = 0.002). (Fig. 2F). For the NMD-sensitive CE, NMD inhibition restored the correlation of CEs and annotated isoforms, while little effect was observed for the non-NMD-rescued CEs (ρ = 0.50, p < 0.001) (Fig. 2G). Overall, localisation of the annotated isoform and sensitivity to NMD drove subcellular localisation of CE.

Finally, we assessed the detection frequency (CE detected in at least two unique samples in a postmortem dataset) and disease selectivity (FPR ≤ 10%, TPR ≥10 % and detected in at least five distinct tissue samples) of NMD-sensitive and NMD-insensitive CEs in brain tissue, and observed that both categories were detected at comparable rates (Fig. 2H), but NMD-insensitive CEs, which include CEs that lead to the production of cryptic peptides^13,14,27^, had greater selectivity for TDP-43 proteinopathy (Fig. 2I).

Finally, while we identified a subset of “dark CE” that only appeared under NMD inhibition in these experiments, we found that most of these events corresponded to previously described CEs (detected in TDP-43 KD datasets lacking NMD inhibition) revealing that “true dark” events are rare and seldom specific to TDP-43 proteinopathy (Fig. 2J; Extended Data Fig. 2E; Supplementary Table 2). For example, the *ATXN1* CE, which appeared as a “dark CE” in previous i^3^ cortical neuron datasets under NMD inhibition^10^ was readily detected in our i^3^ LMNs without NMD inhibition (Fig. 1C). We therefore conclude that while NMD can obscure many CEs in individual experiments, CEs emerging solely upon NMD inhibition are relatively rare, with only 21 “true dark” CE junctions selectively expressed in TDP-43 proteinopathies.

### TDP-43 knockdown causes dose-dependent gene expression and splicing changes

Although *in vitro* TDP-43 KD datasets and postmortem TDP-43 proteinopathy data clearly outline the association of TDP-43 loss and CEs, the levels of TDP-43 loss required for CEs to occur, and whether CEs respond linearly and homogeneously to TDP-43 loss has not yet been addressed. We therefore performed dose-response experiments where clonal SH-SY5Y and SK-N-BE(2) cells were treated with increasing doses of doxycycline to induce progressively stronger TDP-43 KD. We verified TDP-43 levels by qPCR and performed bulk RNA-seq (Fig. 3A; Extended Data Fig. 3A). RNA-seq confirmed the progressive reduction of TDP-43 RNA and also showed that increasing levels of TDP-43 KD led to increased differential gene expression in both cell lines (Fig. 3B).

**Figure 3.**
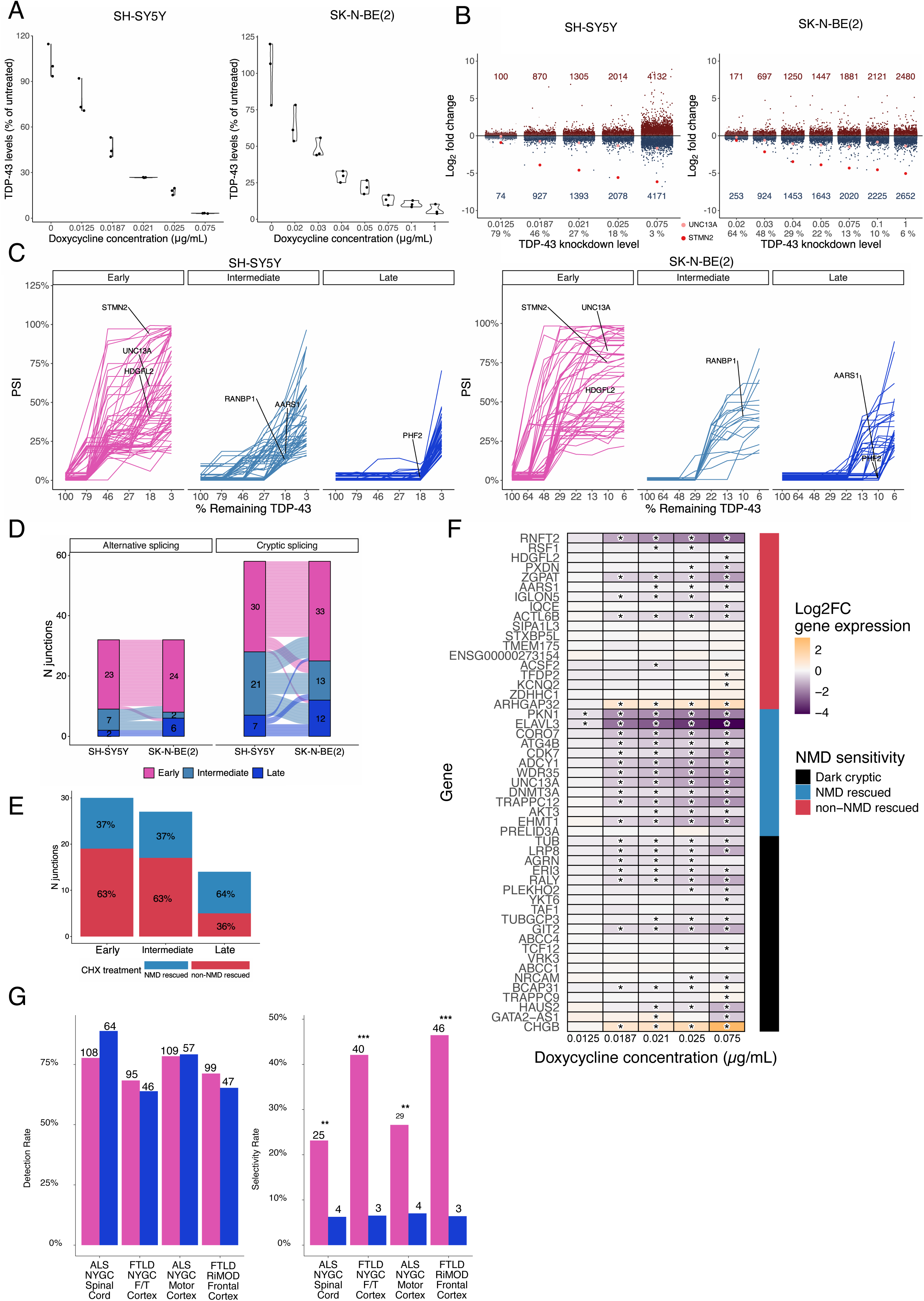
Cryptic splicing shows differential responsiveness to TDP-43 loss of function. A. RT-qPCR in SH-SY5Y (left) and SK-N-BE(2) (right) cells showing levels of TARDBP, normalised to GAPDH, compared to the average in the untreated samples. B. Differential gene expression in the SH-SY5Y (left) and SK-N-BE(2) (right) experiment. Number of up and downregulated genes (adj. p- value < 0.05) called by DESeq2 shown at each level. UNC13A. (salmon) and STMN2 (red) and gene expression highlighted. C. Spaghetti plot showing the response of cryptic splicing events (CE) to TDP-43 loss of function in the SH-SY5Y (left) and SK-N-BE(2) (right) cells. D. Alluvial plot showing the overlap of the early, intermediate, and late-responsive CEs in SH-SY5Y and SK-N-BE(2) cells. E. Number of CE junctions detected in each dose-response category split by NMD- (blue) and non-NMD (red) rescue in SH-SY5Y cells treated with CHX. No significant change in proportion of NMD-rescued vs non-rescued CE, pairwise fisher’s exact test. F. Heatmap visualisation of differential expression of CE-bearing genes categorised by NMD sensitivity in SH-SY5Y cells. Significantly differentially expressed (adj. p-value < 0.05) genes marked with *. G. Detection rate (left) and selectivity rate (right) in the NYGC ALS motor cortex, NYGC ALS spinal cord, NYGC FTLD frontal/temporal cortex, and RiMOD FTLD frontal cortex between early-responsive (pink) and late-responsive (blue) CEs. Numbers above graphs represent the number of CEs meeting either detection or selectivity criteria. Fisher’s exact test. Significance levels reported as * (p <0.05) ** (p <0.01) *** (p<0.001) **** (p<0.0001).

Based on the relative inclusion of CEs at each level of TDP-43 KD, we determined three levels of responsiveness to TDP-43 loss, namely “early”, “intermediate”, and “late” responsive CEs (Fig. 3C). To exclude library depth as a confounding factor in CE detection, we confirmed that sequencing depth did not differ significantly across KD levels (Extended Data Fig. 3B). Genes harboring early-responsive CEs show downregulation at mild TDP-43 KD in both SH-SY5Y and SK-N-BE(2) cells, whereas late-responsive CE genes are only downregulated upon stronger KD (Extended Data Fig. 3C).

TDP-43 depletion also regulates cryptic polyadenylation^18–20^. Therefore, we also assessed the sensitivity of cryptic APA events to differential TDP-43 loss. Using a published list of TDP-43 APAs ^19^ we were able to describe patterns of sensitivity to TDP-43 loss in our progressive TDP-43 KD experiment (Extended Data Fig. 3D). Curiously, we found many more late than early-responsive APA events in the SH-SY5Y dataset.

Cryptic splicing events (PSI < 5% in controls) showed stronger cross-cell line concordance and more consistent TDP-43 responsiveness than alternative splicing events (PSI > 5% in controls) (Fig. 3D), and CE PSI values at maximal TDP-43 depletion were highly correlated between experiments (R = 0.83, p < 0.001) (Extended Data Fig. 3E). Given this strong agreement, we focussed on CEs and combined early and late responsive events across both cell lines into a single category to increase event counts per group.

We then asked if early-responsiveness was due to CEs being less sensitive to NMD degradation. However, no significant difference was seen between the three categories with regard to their NMD status (Fig. 3E;Extended Data Fig. 3F). We next analysed the effect of CE-NMD sensitivity on the responsiveness of parental gene expression to the level of TDP-43 KD in SH-SY5Y cells, restricting this to CEs whose NMD-sensitivity classifications were consistent across both SH-SY5Y NMD-inhibition datasets. Most of the genes harbouring NMD-inducing CEs, such as *ELAVL3* and *UNC13A,* and a set of the “dark cryptics”, including *LRP8* and *GIT2*, exhibited downregulation coherent with TDP-43 KD, whilst the majority of genes with in-frame CEs, such as *HDGFL2* and *KCNQ2*, showed stable or increased expression (Fig. 3F).

Given that CE responsiveness could serve as a potential indicator for staging post-mortem tissue, we next examined the detection and disease selectivity of early and late-responsive cryptic events across post-mortem datasets. Across all datasets, early and late-responsive events were detected at similar rates; however, early-responsive events were much more likely to be selective to TDP-43 proteinopathy (Fig. 3G), and only one late-responsive CE, a cryptic cassette exon in the *AKT3* gene, was consistently selective across all four datasets (Supplementary Table 2).

### Sensitivity of cryptic splicing to TDP-43 loss depends on their features

We set out to understand the sequence features driving the responsiveness of CEs to TDP-43 depletion. We used the brain-specific deep-learning model from Pangolin ^33^ to score the splice site probability of cryptic early and late-responsive splicing events and found that early-responsive cryptic splice sites showed a higher intrinsic splice site probability than late-responsive cryptic splice sites, suggesting that stronger splice sites may need higher levels of TDP-43 for effective repression (Fig. 4A).

**Figure 4.**
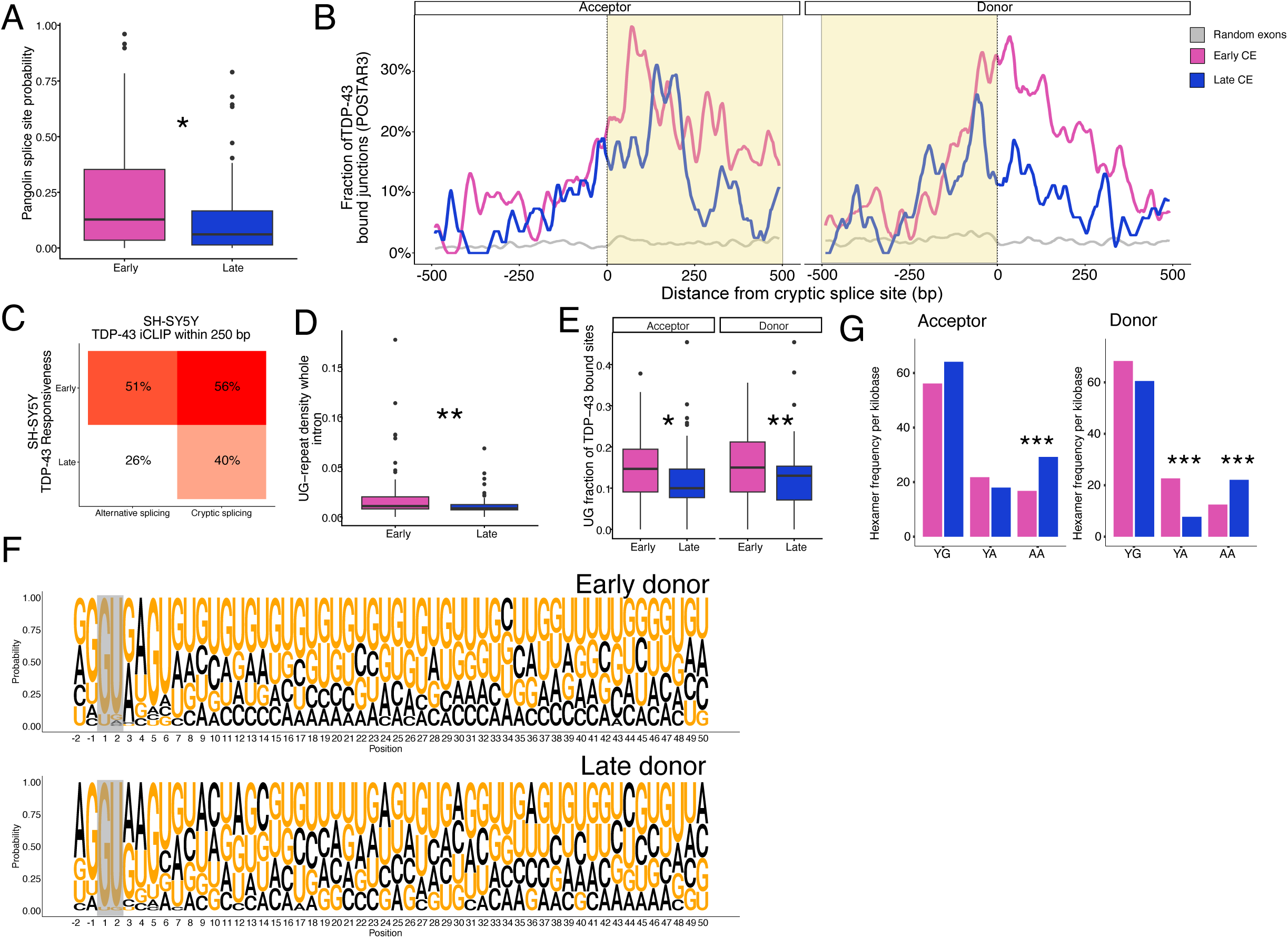
Splice site strength and TDP-43 binding density determine cryptic splicing sensitivity. A. Pangolin splice site probability for early-responsive (pink) and late-responsive (blue) cryptic splicing events (CE). Wilcoxon test. B. Fraction of TDP-43 bound junctions (based on POSTAR3 dataset) around CEs acceptor (left) and donor (right), for early-responsive (pink) and late-responsive (blue) CEs compared to 500 randomly sampled exons (grey). Region inside CEs highlighted in yellow. C. Percentage of TDP-43 iCLIP within 250 base pairs (bp) based on SH-SY5Y iCLIP data between early-responsive and late-responsive cryptic and annotated splicing. D. Whole-intron UG repeat density for early-responsive (pink) and late-responsive (blue) CEs. Wilcoxon test. E. UG fraction of TDP-43 binding sites near acceptor and donor splice sites between early-responsive (pink) and late-responsive (blue) CEs. Wilcoxon test. F. Sequence logo plot depicting probability of each nucleotide for 50 positions from the donor splice site between early-responsive (top) and late-responsive (bottom) CEs based on SH-SY5Y categories. Splice site highlighted in grey. G. Hexamer frequency of TDP-43 bound regions by TDP-43 binding category36 per kilobase in acceptor (left) and donor (right) splice site between early-responsive (pink) and late-responsive (blue) CEs. Fisher’s exact test. Box plots: boundaries 25–75th percentiles; midline, median; whiskers, Tukey style. Significance levels reported as * (p <0.05) ** (p <0.01) *** (p<0.001) **** (p<0.0001).

Next, we analyzed TDP-43 binding patterns around these splice sites. Using the POSTAR3 database ^34^, which aggregates multiple TDP-43 crosslinking and immunoprecipitation (CLIP) studies, we calculated the fraction of junctions with a TDP-43 binding site within +/- 500 basepairs of the cryptic splice sites. Compared to randomly sampled exons, both early and late-responsive CEs had an enrichment of TDP-43 binding downstream of the acceptor site within the cryptic exon and around the donor site. In both instances early-responsive events show a higher enrichment compared to late-responsive events (Fig. 4B).

To confirm this, we used TDP-43 iCLIP from SH-SY5Y cells ^8^, where also the dose-response experiment was performed, and observed early-responsive splicing events, both cryptic (PSI < 5% in controls) and alternative (PSI > 5% in controls), were more likely than late-responsive splicing events to have a TDP-43 binding site within 250 basepairs of the splice site (Fig. 4C). Moreover, introns with early-responsive CEs had a higher UG-repeat density than late-responsive CEs across the entire intron, consistent with classic TDP-43 UG binding action (Fig. 4D).

To further characterize TDP-43 binding in these introns, we analysed the sequences where TDP-43 binding had been observed by CLIP, and found TDP-43 bound regions of early-responsive CEs had higher UG-frequency (Fig. 4E). A sequence logo analysis^35^ of the first 50 base pairs of the intron downstream of the donor site revealed early-responsive donor sequences were dominated by uninterrupted UG repeats, whereas late-responsive donor sequences, although UG-rich, contained more interruptions (Fig. 4F).

Early-responsive CEs required lower levels of TDP-43 depletion to occur yet appeared to contain both more TDP-43 binding and more UG density. TDP-43’s RNA binding is tuned by its ability to phase-separate and form RBP condensates, with condensation-dependent binding preferring longer stretches of canonical YG-containing and YA-containing [UG]n motifs, and condensation-independent binding occurring at shorter AA-containing [UG]n motifs, where the repetitive UG pattern is interrupted with an AA dinucleotides^36^. Using these motif classes, we assessed motif enrichment in TDP-43-bound regions by extracting the sequences of POSTAR3 TDP-43 CLIP sites^34^ within +/- 500 basepairs of the cryptic splice sites. Both early and late-responsive CEs most frequently used the canonical YG motifs in binding regions flanking acceptor and donor splice sites (Fig. 4G). However, late-responsive CEs had an enrichment for AA-containing motifs at both acceptor and donor sites (0.80 and 0.82 log_2_fold motif frequency respectively, adj. p-value Fisher’s test < 0.001), and depletion of YA-containing motifs at late-responsive donor sites compared to early-responsive events (-1.56 log_2_fold motif frequency, adj. p-value Fisher’s test < 0.001) (Fig. 4G). The distribution of motif classes and binding profiles resembles the condensation-dependent versus-independent TDP-43 binding modes described by Hallegger, with early-responsive events aligning more closely to the condensation-dependent pattern.

Together these results demonstrate that early-responsive cryptic events occur with stronger splice sites and show more UG-rich TDP-43 binding motifs with longer TDP-43 binding regions, features similar to condensation-dependent binding. These factors work together to likely make them more sensitive to loss of TDP-43 repression.

### A composite cryptic burden score captures disease heterogeneity

In our postmortem datasets, we next asked if we could quantify the overall load of CE inclusion per sample. To do this we selected CEs that met predictive criteria (FPR ≤ 10%, TPR ≥10 % and detected in at least five distinct TDP-43 proteinopathy tissue samples) in each dataset and constructed a combined cryptic burden score by summing the dataset normalised CE PSI.

We then compared the ability of the cryptic burden score to distinguish TDP-43 proteinopathy. In the two FTD datasets, the cryptic burden score performed comparably to the strongest single-event predictors (RiMOD: Area under curve (AUC) = 1.0; *HS6ST3* = 0.998; *KCNQ2* = 0.994; NYGC: AUC = 0.967; *STMN2* = 0.934; *KCNQ2* = 0.934) (data not shown). However, in the ALS datasets, especially in the motor cortex, the cryptic burden score outperformed the top individual CE predictors (Fig. 5A). These findings suggest that aggregating cryptic events into a composite score may offer a more reliable indicator of TDP-43 proteinopathy, especially in heterogeneous ALS tissues where individual markers may show limited detection.

**Figure 5.**
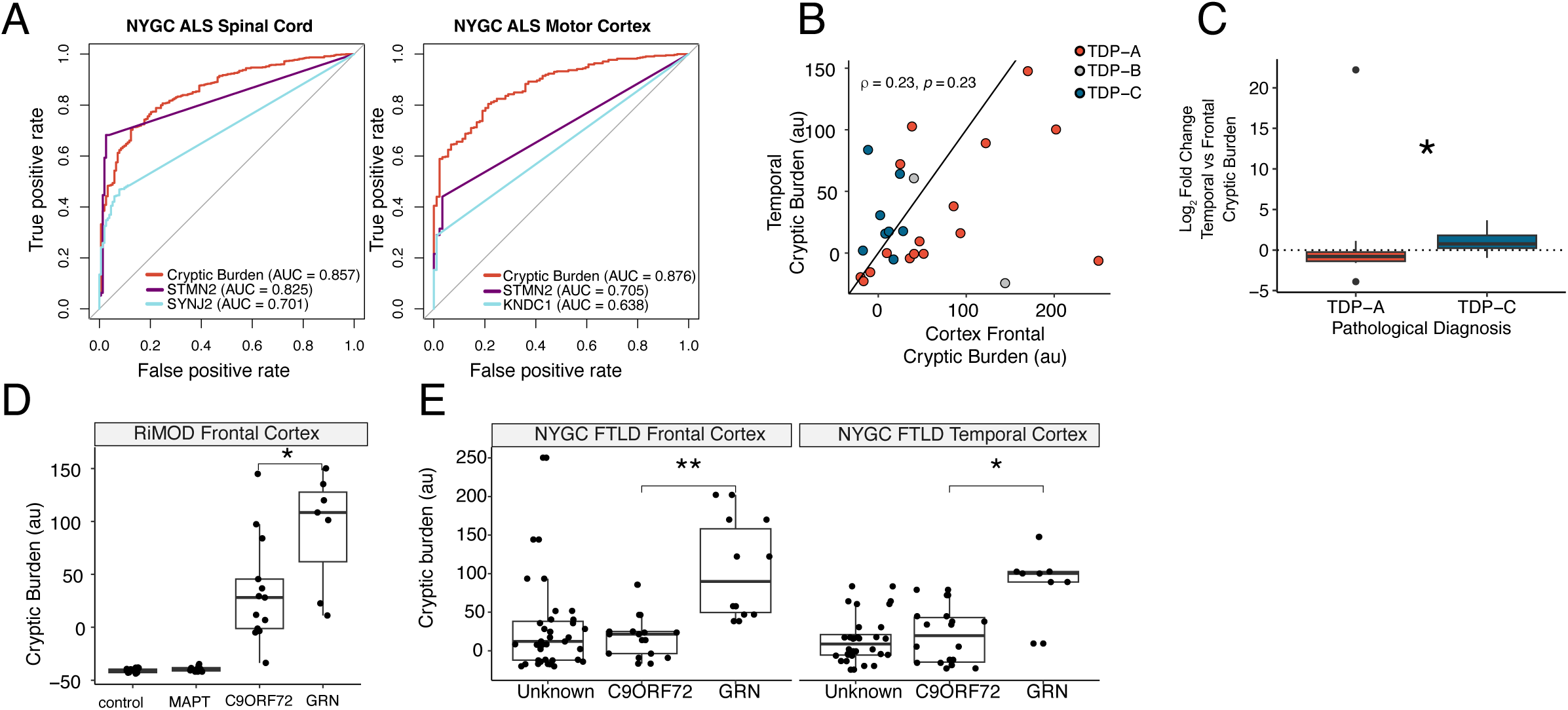
A cryptic burden score reveals disease heterogeneity. A. Receiving operator curves using cryptic splicing burden or top two individual CE predictors in NYGC ALS spinal cord (left) and NYGC ALS motor cortex (right). B. Scatter plot displaying frontal cortex (x-axis) and temporal cortex (y-axis) cryptic exon burden in NYGC FTLD cases split by pathological type (A, B, or C). C. Differential cryptic splicing burden between temporal and frontal cortex in the same patient for NYGC FTLD type A and type C. Wilcoxon test. D. Cryptic splicing burden between controls and different genetic types of FTLD (MAPT, C9orf72, and GRN) in the RiMOD dataset. Wilcoxon test. E. Cryptic splicing burden between different genetic types of FTLD (unknown, C9orf72, and GRN) in NYGC FTLD frontal cortex (left) and NYGC FTLD temporal cortex (right). Wilcoxon test. Box plots: boundaries 25–75th percentiles; midline, median; whiskers, Tukey style. Significance levels reported as * (p < 0.05) ** (p < 0.01) *** (p<0.001) **** (p<0.0001).

Across spinal cord ALS-TDP samples, the lumbar region exhibited the highest cryptic burden compared to thoracic or cervical regions (Extended Data Fig. 4A), and cryptic burden in the motor cortex showed a low but significant correlation with lumbar cord (ρ = 0.15, p = 0.017) from the same ALS-TDP cases (Extended Data Fig. 4B). In the NYGC frontal and temporal cortices, there was no significant correlation between cryptic burden between the frontal and temporal cortex (Fig. 5B). However, when we stratified by TDP-43 type, consistent with known regional pathology ^37,38^, type C TDP-43 cases showed greater cryptic load in the temporal cortex, whereas type A cases had higher cryptic burden in frontal cortex (Fig. 5C).

We next examined whether genetic background influenced cryptic burden. In the RiMOD dataset, *GRN* mutation carriers displayed higher cryptic burden than *C9orf72* carriers (Fig. 5D). *GRN* mutations are typically associated with TDP-43 type A^39^. Similarly, in the NYGC cohort, *GRN* carriers showed higher cryptic burden in both frontal and temporal cortices, despite type A pathology generally having a higher cryptic burden in the frontal cortex in our data (Fig. 5E). To assess whether this difference reflected mutation status rather than pathology type, we restricted the analysis in the NYGC datasets to confirmed type A cases. Within this subset, *GRN* carriers still showed a trend toward higher cryptic burden, although the difference was no longer statistically significant (Extended Data Fig. 4C). Finally, to aid researchers in exploring the consequences of the NMD-sensitivity, specificity, detection of all cryptic events analysed, and gene expression in response to progressive TDP-43 depletion we provide a web-based browser at: https://anna-leigh-brown.shinyapps.io/Cryptic-Creeper/

## Discussion

Over the last decade, cryptic splicing triggered by TDP-43 loss of function has emerged as a fundamental pathogenic mechanism in neurodegenerative diseases including ALS, FTD and AD. Research has focused on a subset of these events, for either their genetic links to ALS/FTD (*UNC13A*), compelling roles in ALS pathogenesis (*STMN2*, *KCNQ2*, *ATG4B* and *G3BP1*), or utility as disease biomarkers (*HDGFL2*)^8,9,13,14,16,17,26,27,40–42^. Although these CEs along with others have been analysed in *post mortem* brains, a comprehensive transcriptome-wide view of CEs in disease and how their biological features, such as degradation susceptibility, subcellular localisation, and sensitivity to TDP-43 loss, relate to pathology remains lacking. Our work addresses these gaps.

TDP-43 aggregation and low levels of mis-splicing has been reported in healthy aging brain^43^. While some cryptic events like *EP400*^23^ may be enriched in disease, we focused on identifying disease selective events, which are more likely to yield specific biomarkers or therapeutic targets. While we discovered over 5,000 unique cryptic junctions across all knockdowns *in vitro*, only 5% were specific to TDP-43 proteinopathy in diseased brains. This poor discrepancy may reflect limited cell diversity *in vitro*. For example, the cryptic skipping event in *KCNQ2* is specific to TDP-43 proteinopathy in brain tissue, but is highly expressed in normal male reproductive tissue^23^, implying some “cryptic” events in neurons may be normal but unannotated in other cell types.

Defining particular biological features of individual cryptic splicing events, such as their potential to produce cryptic peptides^13,14,27^, subcellular localization, and sensitivity to TDP-43 depletion is crucial for biomarker development. Most cryptic splicing events are known to lead to gene loss through NMD^7,12^. We and other groups^8,10,11^ have reported that NMD can mask the detection of cryptic exons. Yet by combining our three orthogonal approaches to NMD inhibition with our analyses of TDP-43 knockdown datasets, we discovered that ‘true dark’ cryptics, detectable only under both TDP-43 knockdown and NMD inhibition, are rare. Most dark CEs from one dataset appeared in other TDP-43 knockdown datasets without NMD inhibition, suggesting cell-type-specific NMD efficiency. Furthermore, we found no evidence of a general NMD deficiency following TDP-43 knockdown. Combining NMD inhibition experiments with our knockdown datasets, we identified 497 ‘true dark’ events, with 21 being detectable and specific to TDP-43 proteinopathy brains.

As expected, NMD depleted CE RNA from cytoplasmic fraction, but by inhibiting NMD, thus removing its cytoplasmic depletion, we found that most CE-containing transcripts localised similarly to their canonical isoforms. Interestingly some CEs, like *RAP1GAP*, maintained nuclear enrichment despite NMD inhibition, suggesting there may be additional RNA regulatory measures controlling their nuclear detention. Detecting CEs in RNA from brain-derived extracellular vesicles (EVs) is a promising non-invasive biomarker for diseases, however RNA must be both stable and cytoplasmic to be loaded into EVs^44^. Thus, our approach identified a cohort of CEs that are not only NMD-stable, but also cytoplasmically-localised and specific to TDP-43 proteinopathy, marking them as prime candidates for future studies into EV-derived RNA.

Although the detection of CEs indicates TDP-43 loss in diseased tissue, TDP-43 nuclear depletion occurs at varying degrees, and how CEs relate to TDP-43 levels is still largely unknown. Our datasets which include RNA-seq on 12 different levels of TDP-43 expression across two different neuronal lines represent a unique resource. Our data enable development of probe-based assays to visualise graded TDP-43 dysfunction and to define the order of CE emergence in postmortem brain samples and longitudinal biofluids. Understanding CE sensitivity to TDP-43 loss is also important for understanding disease pathogenesis. For example, we find that *KCNQ2* and *UNC13A* are both early events, but *KCNQ2* appears before *UNC13A*. Whilst cryptic *KCNQ2* induces neuronal hyperexcitability^26^, cryptic *UNC13A* dampens synaptic transmission^24^, and the order in which these two mis-splicing events appear may underpin the different phases of neuronal hyperexcitability and hypoexcitability observed in disease. Multiple other factors in trans, such as the levels of other RBPs, likely modulate TDP-43 responsiveness. Recent studies showing that individual cryptic exons can be co-regulated by TDP-43 and distinct RBPs, e.g. *G3BP1*-CE and SRRM4, *HDGFL2*-CE and SRSF3 and *UNC13A*-CE and hnRNPL^29,41,45,46^, suggest that cryptic events may be individually modulated by a unique set of regulatory factors. That early-responsive events were also more specific in post mortem brain datasets is consistent with early-responsive CEs being under a more direct TDP-43 regulation, whereas late events may also reflect downstream or secondary cellular changes.

By integrating *in vivo* predictive CEs, our composite TDP-43 cryptic burden score captures TDP-43 dysfunction more sensitively than single targets. This score reflected the expected regional patterns of TDP-43 pathology, correlating with predominant frontal lobe involvement in FTD type A and temporal lobe involvement in type C. It also revealed a higher cryptic burden in FTD-GRN compared to FTD-C9orf72. Neuronal and glial cytoplasmic pTDP-43 inclusions are equally severe in FTD-GRN and FTD-C9orf72, but FTD-GRN is associated with more pro-inflammatory reactive microglia^47,48^. Given the increased cryptic burden in FTD-GRN, a critical question for future research may be the relationship between cryptic splicing burden and inflammatory microglia.

Despite the advances of this work, several limitations remain. Functional validation is needed to clarify how CE inclusion contributes to protein dysfunction, cellular toxicity, and disease mechanisms. Our analyses were restricted to specific models and brain regions and extending to additional cell types, tissues, and diseases will test the generality of these findings. Finally, because standard 10x single-cell datasets detect primarily 3’ CEs with low sensitivity^23,49^, we excluded them from our analyses. Future probe-based and full-coverage single-cell approaches will better resolve CE timing, co-occurrence, and cell-type specificity.

Our work opens several avenues for further study. Prioritising key CEs for mechanistic and functional validation will clarify their pathogenic roles. Expanding analyses of TDP-43 loss-of-function splicing across diverse cell types, tissues, and diseases may reveal shared and context-specific RNA processing defects. Finally, combining multiple CEs into biomarker panels could improve sensitivity and specificity for detecting and subtyping TDP-43 proteinopathies, advancing diagnosis and disease monitoring.

To aid the field we provide an accompanying Shiny app that allows open access to explore cryptic splicing across in vitro and brain datasets. This resource is designed to incorporate future data from additional cell types, maintaining an up-to-date reference for the field.

## Tables

Supplementary Table 1. Gene expression changes in all TDP-43 knockdown datasets.

Supplementary Table 2. Summary of all detected cryptic including postmortem detection, TDP-43 sensitivity, and NMD sensitivity.

Supplementary Table 3. Summary of samples from NYGC and RiMOD cohorts.

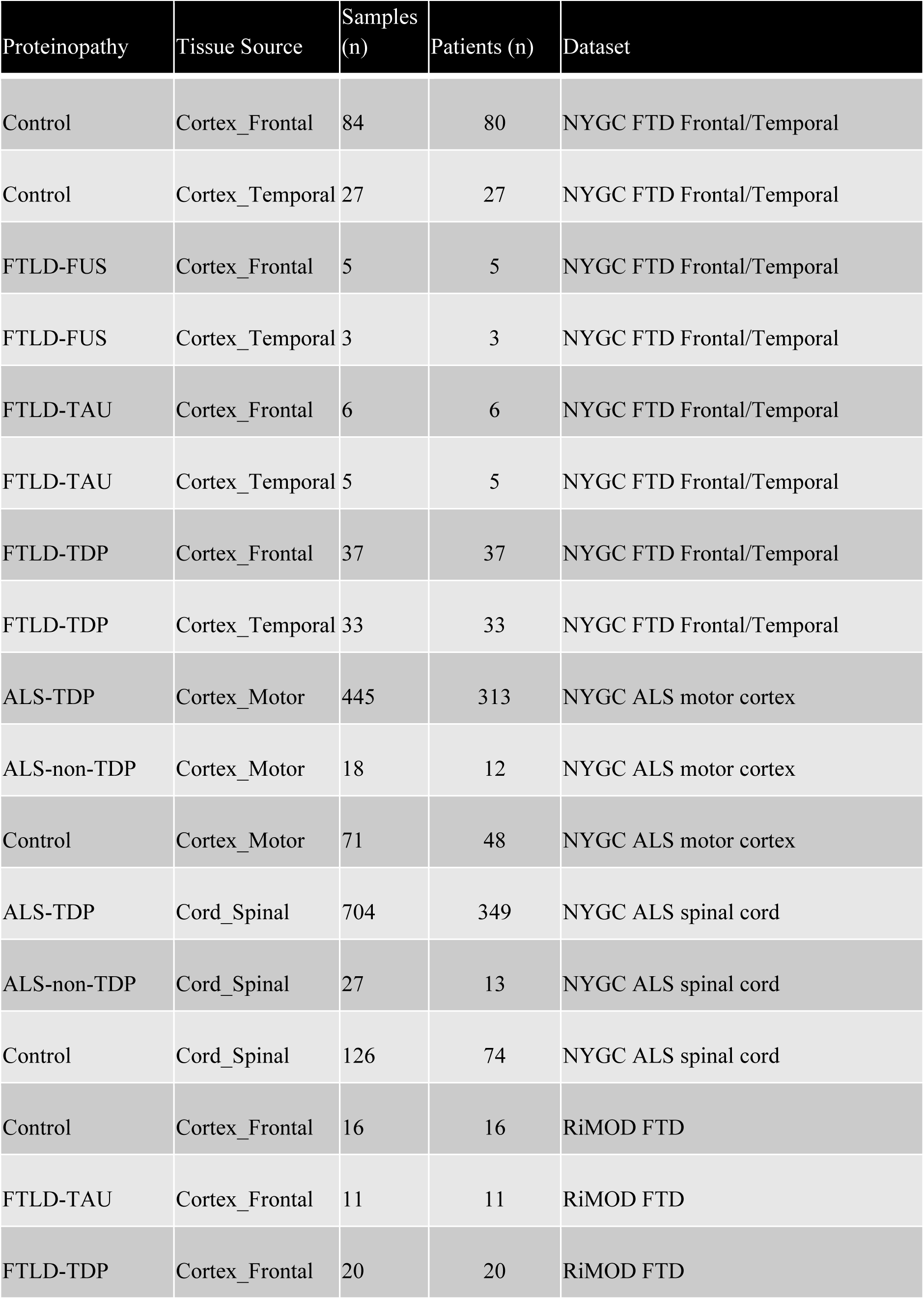

## Methods

### Previously published data

Publicly available data were obtained from the Gene Expression Omnibus (GEO):

– iPS cell MNs ‘controlklimmotorneuron-tdp43kdklimmotorneuron’^16^, GSE121569;
– - SK-N-BE(2)^50^, GSE97262;
– FACS-sorted frontal cortex neuronal nuclei^31^, GSE126543;
– Controlfergusonhela-tdp43kdfergusonhela^51^
– Controlbrowncorticalneuron-tdp43kdbrowncorticalneuron^8^
– Controlbrownshsy5y-tdp43kdbrownshsy5y^8^
– Controlbrownskndz-tdp43kdbrownskndz^8^
– Controlhumphreycorticalneuron-tdp43kdhumphreycorticalneuron^19^
– Controlmelamedshsy5y-tdp43kdmelamedshsy5y^17^
– Controlseddighicorticalneuron-tdp43kdseddighicorticalneuron^14^

### Human iPSC culture

Human iPS cells used in this study were engineered^52,53^ to express mouse or human neurogenin-2 or hNIL (NGN2, ISL1, LHX3) as well as an enzymatically dead Cas9 (+/− CAG-dCas9-BFP-KRAB)^54^. To achieve knockdown, sgRNAs targeting either a non-targeting control guide (GTCCACCCTTATCTAGGCTA), *UPF1* (GGCCAGACGCAGACGCCCCC), or *TARDBP* (GGGAAGTCAGCCGTGAGACC) were delivered to iPS cells by lentiviral transduction^55^. For NMD inhibition experiments, the following conditions were used:

– Control for NMD and control for TDP-43 knockdown (control/control)
– Positive for UPF1 knockdown and control for TDP-43 knockdown(UPF1KD/control)
– Control for NMD and positive for TDP-43 knockdown and c (control/TDP43KD)
– Positive for UPF1 knockdown and positive for TDP-43 knockdown(UPF1KD/TDP43KD) For other experiments, cells were transduced only with control or *TARDBP*-targeting sgRNA.

For i3cortical-like neurons, after lentiviral transduction cells were cultured in tissue culture dishes coated with human embryonic stem cell-qualified Matrigel (Corning). The following morning, cells were washed with PBS, and the media was changed to E8 alone or (depending on cell number) E8 supplemented with 10 µM ROCK inhibitor in a 37°C, 5% CO2 incubator. Cells were passaged with accutase (Life Technologies) for 5–10 min at 37 °C and then washed with PBS before re-plating. Two days after lentiviral delivery, cells were selected overnight with either puromycin (10 µg/ml) or blasticidin (50–100 µg/ml). iPS cells were then expanded for 2 days before initiating neuronal differentiation. To initiate neuronal differentiation, 20–25 million iPS cells per 15 cm plate were individualised using accutase on day 0 and re-plated onto Matrigel-coated tissue culture dishes in N2 differentiation media containing: knockout DMEM/F12 media (Life Technologies Corporation) with N2 supplement (Life Technologies Corporation), 1× GlutaMAX (Thermofisher Scientific), 1× MEM nonessential amino acids (Thermofisher Scientific), 10 µM ROCK inhibitor (Selleckchem) and 2 µg/ml doxycycline (Clontech). The media was changed daily during this stage. On day 3, pre-neuron cells were replated onto dishes coated with freshly made poly-L-ornithine (0.1 mg/ml; Sigma), 6-well dishes (2 million per well), in i3Neuron Culture Media: Brain-Phys media (Stemcell Technologies) supplemented with 1× B27 Plus Supplement (ThermoFisher Scientific), 10 ng/ml BDNF (PeproTech), 10 ng/ml NT-3 (PeproTech), 1 µg/ml mouse laminin (Sigma), and 2 µg/ml doxycycline (Clontech). i^3^ Neurons were then fed three times a week by half media changes. i^3^ Neurons were then collected on day 7 after the addition of doxycycline.

For i^3^ LMNs, iPSCs were transduced in suspension after dissociation with Stem Pro Accutase (Thermo Fisher), plated in E8 Flex media supplemented with 10 µM ROCK inhibitor (Y-27632, Tocris Bioscience) and cultured overnight in a 37°C, 5% CO2 incubator. Cells were then washed with PBS and media was changed to E8 Flex media. After 48 hours from transduction, cells were passaged with Stem Pro Accutase and replated in E8 Flex media supplemented with 10 µM ROCK inhibitor and puromycin (10 µg/ml) for selection. Before differentiation, cells were expanded for 2 days in E8 Flex media. To induce neuronal differentiation, induction medium (DMEM/F12, GlutaMAX supplement medium (Gibco, 31331028), MEM non-essential amino acids (Gibco, 1140050), N2 Max supplement (R&D Systems, AR009), 0.2 µM compound E, 2 μg/ml doxycycline hyclate (Sigma-Aldrich, D9891)) was added to the cells on day 0. After 48 hours of induction, cells were re-plated onto dishes coated with 50 μg/ml Poly-D-Lysine (Gibco, A38904-01) and 10 μg/ml laminin (Gibco, 23017015) in induction medium supplemented with 10 μM ROCK inhibitor, 1 μg/ml laminin and CultureOne supplement (Gibco, A3320201). After 24 hours (day 3), induction medium was replaced with motor neuron medium containing Neurobasal (Gibco, 21103049), MEM non-essential amino acids, GlutaMAX supplement (35050038), N2 max supplement (R&D Systems, AR009), N21 max supplement (R&D Systems, AR008), CultureOne supplement (Gibco, A3320201), 1 μg/ml Laminin, 2 μg/ml doxycycline hyclate, 10 ng/ml BDNF (Peprotech) and 10 ng/ml GDNF (Peprotech). Half-medium changes were performed twice a week. Cells were maintained in an incubator at 37°C and 5% CO2 and collected on day 7.

### Cell lines with dox-inducible TDP-43 knockdown

SH-SY5Y and SK-N-BE(2) cells were transduced with SmartVector lentivirus (V3IHSHEG_6494503) containing a shRNA cassette for TARDBP, which can be induced by doxycycline. Transduced cells were selected with puromycin (1 µg/ml) for one week. The pool of TDP-43-knockdown SH-SY5Y and SK-N-BE(2) cells was plated as single cells and expanded to obtain a clonal population. Cells were grown in DMEM/F12 containing Glutamax (Thermo) supplemented with 10% FBS (Thermo) and 1% PenStrep (Thermo). For induction of shRNA against TARDBP, cells were treated for 10 days with increasing amounts of doxycyline hyclate (Sigma), as follows:

– For experiments in SH-SY5Y cells, 12.5 ng/ml, 18.75 ng/ml, 21 ng/ml, 25 ng/ml, and 75 ng/ml.
– For experiments in SK-N-BE(2) cells, 20 ng/ml, 30 ng/ml, 40 ng/ml, 50 ng/ml, 75 ng/ml, 100 ng/ml, and 1000 ng/ml.

### NMD inhibition in SH-SY5Y cells

For the NMD inhibition experiment with cycloheximide (CHX) in SH-SY5Y, 10 days after the induction of shRNA against TDP-43 with 25 ng/ml doxycyline hyclate (Sigma), cells were treated either with 100 µM CHX or DMSO^56^ for 6 h before isolating the RNA. Overall, 4 conditions were generated:

– Control for TDP-43 knockdown and control for NMD (control/control)
– Control for TDP-43 knockdown and CHX-treated (control/CHX)
– Positive for TDP-43 knockdown and control for NMD (TDP43/control)
– Positive for TDP-43 knockdown and CHX-treated (TDP43/CHX)

### Cell fractionation experiment

For the fractionation experiments, SH-SY5Y cells were treated for 10 days with 25 ng/ml doxycycline hyclate (Sigma) to induce the shRNA against TDP-43, and for 24 hours with 0.5 μM SMG1-11j inhibitor^57^ to block the NMD machinery. After 10 days, cells were trypsinised, pelleted and resuspended in 1X PBS (Thermo). A fraction of the resuspended cells was pelleted and used for the subcellular fractionation with the Ambion PARIS Kit (Life Technologies), according to the manufacturer’s instructions. RNA from the nuclear and the cytosolic fractions was extracted with the Direct-zol kit (Zymo) with on-column DNase I treatment. For each experimental condition, 2 μg of cytoplasmic RNA and an equal volume of nuclear RNA fraction were reverse-transcribed with RevertAid First Strand cDNA Synthesis Kit (Thermo Fisher Scientific) according to the manufacturer’s instructions.

### RNA sequencing, differential expression and splicing analysis

Sequencing libraries were prepared with polyA enrichment using a TruSeq Stranded mRNA Prep Kit (Illumina) and sequenced on an Illumina HiSeq 2500 or NovaSeq 6000 machine at UCL Genomics with the following specifics:

– SH-SY5Y cycloheximide and dose-response curves: 2 × 100 bp, depth > 40M/sample
– SK-N-BE(2) dose-response curves, SH-SY5Y fractionation experiment, and UPF1 iPSCs: 2x150 bp, depth > 40M/sample

Raw sequences (in fastq format) were trimmed using Fastp^58^ with the parameter “qualified_quality_phred: 10”, and aligned using STAR (v2.7.0f)^59^ to the GRCh38 with gene models from GENCODE v40^60^. The STAR outputs are BAM files, tab-delimited files containing the counts and coordinates for all splice junctions found in the sample, and alignment metadata including the genomic location where a read is mapped to, read sequence, and quality score. Then, using samtools^61^, BAM files were sorted and indexed by the read coordinates location. Trimmed fastqc files were aligned to the transcriptome using Salmon (v1.5.1)^62^, outputting isoform-specific counts used for differential gene expression, performed using DeSEQ2^63^ without covariates, using an index built from GENCODE v40^60^. The DESeq2 median of ratios, which controls for both sequencing depth and RNA composition, was used to normalise gene counts. Differential expression was defined at a Benjamini–Hochberg false discovery rate < 0.1. We kept genes with at least 5 counts per million in more than 2 samples. Splicing analysis was performed using Majiq (v2.1)^64^ on STAR-aligned and sorted BAMs and the GRCh38 reference genomes. MAJIQ’s outputs were used to categorise each of the splicing junctions by the overlap with annotated transcripts and exons, using GTF and DASPER (https://github.com/dzhang32/dasper), classifying them into “annotated”, “novel_acceptor”, “novel_donor”, “novel_exon_skip”, “novel_combo” and “ambig_gene” and “none”. Cryptic splicing was defined as junctions with Ψ (PSI, percent spliced in) < 5% in the control samples, ΔΨ > 10% between groups, and the junctions were unannotated in GENCODE v40^60^. In some cases, for a cryptic exon, there was only enough coverage to call one of the two defining junctions significant. QC reports were generated with FASTQC and then summarised using MultiQC^65^. The alignment and splicing pipelines are implemented in Snakemake (v5.5.4)^66^, a workflow management software that allows reliable, reproducible, and scalable analysis, and runs on the UCL high-performance cluster. PCA analysis and data visualisation were run on R using custom scripts made publicly available.

### RNA-seq data processing

‘Humphrey i3 Cortical’ samples were processed as previously described^19^ using the RAPiD-nf nextflow pipeline. Briefly, adapters were trimmed from raw reads using Trimmomatic^67^ v0.36, and reads were aligned to the GRCh38 genome build using gene models from GENCODE v30^60^ with STAR^59^ v2.7.2a. The RAPiD-nf pipeline is available at https://github.com/CommonMindConsortium/RAPiD-nf/. The ‘Brown’ SH-SY-5Y, SK-N-BE(2) and i3Neuron datasets were processed as previously described^8^. Unless otherwise stated, all short-read RNA-seq datasets were processed using the following pipeline. Raw reads in FASTQ format were quality trimmed for a minimum Phred score of 10 and otherwise default parameters using fastp^58^ (v0.20.1). Quality trimmed reads were aligned to the GRCh38 genome using gene models from GENCODE v40^60^ with STAR^59^ (v2.7.8a). Quality trimmed reads are used as input for any tools that require FASTQ files as input (e.g. PAPA, Salmon). Our alignment pipeline is implemented in Snakemake^66^ and is available at https://github.com/frattalab/rna_seq_snakemake.

### Classification into “early”, “intermediate” and “late” events

To define “early”, “intermediate”, and “late” categories of alternative splicing/polyadenylation in the TDP-43 knockdown curves, we first visually screened IGV tracks to shortlist *bona fide* alternative splicing events, keeping only events with at least 5 splicing junctions (on both sides for cassette events). We then assigned a CE to the “early” category if its PSI reached 20% in either the first or second TDP-43 knockdown level, and to the “late” category if its PSI reached 10% only upon the strongest level of TDP-43 knockdown. All the other events were binned into the “intermediate” category.

### Motif analysis, GU, splice strength

UG-repeat density was determined as the number of TGTGTG sequences (allowing one mismatch) in a sequence of a canonical intron, normalised to intron length. For TDP-43 iCLIP sites specifically from SH-SY5Y cells we used previously published peaks ^8^. For all TDP-43 binding sites we downloaded the data from POSTAR3 ^34^ and collapsed overlapping binding regions from multiple studies into a single binding region. TDP-43-enriched hexamer motifs were analyzed based on previously characterized binding sequences^36^, grouped into three categories: YG (TGTGTG, GTGTGT, TGTGCG, TGCGTG, CGTGTG, GTGTGC), YA (ATGTGT, GTATGT, GTGTAT, TGTGTA, TGTATG, TGCATG), and AA (GTGTGA, AATGAA, GAATGA, TGAATG, ATGAAT, GTGAAT, GAATGT, TTGAAT). Genomic sequences were extracted from splice junction flanking regions that overlapped with TDP-43 binding sites.

Motif frequencies were calculated as occurrences per kilobase for each hexamer group. Enrichment of motifs in late versus early junctions was assessed using Fisher’s exact test on 2×2 contingency tables (motif present vs. absent), with Benjamini-Hochberg correction for multiple testing. Results with adjusted p-values < 0.05 were considered statistically significant, with fold-enrichment calculated as the ratio of motif frequencies between conditions. Sequence logo visualisation was performed in R 4.3.2, using the ggseqlogo package ^35^. For Pangolin ^33^ splice site strength prediction we extracted the genomic sequence 5000 basepairs up and downstream of the cryptic splice sites and ran splice site prediction using the brain specific models. Code to run Pangolin splicing prediction is available at https://github.com/aleighbrown/pangolin_run_cryptic_biology/blob/main/PangolinColab.ipynb.

### Cryptic last exons definition

To assess the levels of inclusion of TDP-43 cryptic last exons in our doxycycline-titration experiment, we employed the PAPA pipeline recently published by our group^19^ Briefly, we ran PAPA (https://github.com/frattalab/PAPA) in ‘identification’ mode to predict novel last exons in the doxyycline-titration SH-SY5Y and SK-N-BE(2) datasets. We provided GENCODE v42 annotations filtered for protein-coding and lncRNA gene transcripts with a ‘transcript support level’ value≤3 and without the ‘mRNA_end_NF’ tag. Predicted last exon GTF files were combined into a single GTF using PAPA’s ‘combine_novel_last_exons.py’ script. All datasets were then quantified and assessed for differential usage using a unified transcriptome reference combining novel and annotated last exons from the filtered GTF. Differential usage was performed using the standard DEXSeq workflow. We defined cryptic APAs as DEXSeq adjusted P<0.05, mean control usage<10% and change in mean usage>10% (TDP-43 knockdown, control). We further manually curated cryptic IPAs, as manual inspection suggested frequent artifacts at regions of reduced coverage in intron retention loci. The same classification system for early and late responsive CEs was used as for cryptic last exons events.

### RNA extraction and RT-(q)PCR

RNA was extracted from SH-SY5Y, SK-N-BE(2) with an RNeasy mini kit (Qiagen) following the manufacturer’s protocol, including the on-column DNA digestion step. RNA concentrations were measured by Nanodrop, and 500–1000 ng of RNA was used for reverse transcription. Samples undergoing RNA sequencing were furthermore assessed for RNA quality on a TapeStation 4200 (Agilent), and bands were quantified with TapeStation Systems Software v3.2 (Agilent). The RNA integrity number (RIN) was above 9.4.

Reverse transcription was performed from 1000 ng of RNA using RevertAid cDNA synthesis kit (Thermo) using random hexamer primers and following the manufacturer’s protocol. Transcript levels were quantified by qPCR (QuantStudio 5 Real-Time PCR system, Applied Biosystems) using the ΔΔCt method^68^. Using RefFinder^69^, we identified GAPDH as the most stable endogenous control across our conditions of interest. Gene expression analysis of *TARDBP* was assessed by means of qPCR, using the following primers: *TARDBP* forward, 5′-GATGGTGTGACTGCAAACTTC-3′; *TARDBP* reverse, 5′-CAGCTCATCCTCAGTCATGTC-3′.

### Analyses of bulk post-mortem datasets

We analyzed RNA-seq data from the NYGC ALS cohort and the RiMOD-FTD project using a uniform processing pipeline. Reads were aligned to GRCh38 with the RaPID-nf pipeline (https://github.com/CommonMindConsortium/RAPiD-nf) with STAR v2.7.2a^70^. Library depth was measured as the number of uniquely mapped reads with Picard CollectAlignmentSummaryMetrics. Sample processing, library preparation, and initial RNA-seq QC for both NYGC and RiMOD-FTD cohorts have been described previously^71–73^.

Splice junctions were extracted from aligned BAM files with RegTools^74^ using an 8 bp anchor and 500 kb maximum intron size, then clustered with LeafCutter^75^ under relaxed filtering (≥0.0001 fraction of cluster reads). Gene expression was quantified as TPM using TPMCalculator^76^ and taken from the ‘_genes.out’ output using gene models from GENCODE v40^60^.

### Splicing event classification ability in bulk post-mortem datasets

To assess the predictive capability of individual splicing events for TDP-43 proteinopathy, samples in each bulk post-mortem dataset were assigned a binary classification of either “TDP-43 proteinopathy” or “non-TDP-43 proteinopathy”. For the NYGC ALS cohort, all ALS cases, excluding those with SOD1 or FUS mutations, were categorized as “TDP-43 proteinopathy”, while controls and SOD1/FUS samples were classified as “non-TDP-43 proteinopathy”. TDP-43 pathology was confirmed post-mortem for all FTD samples included. In the RiMOD dataset, GRN and C9orf72 cases were considered “TDP-43 proteinopathy”, and controls and MAPT cases were classified as “non-TDP-43 proteinopathy”. To determine if a splice junction could predict between these two classes, we first z-score normalised the PSI of each junction across all samples in that specific dataset to ensure comparability across events with different expression ranges. For each normalized event, we constructed receiver operating characteristic curves using the pROC R package^77^ calculated the area under the curve, along with the true and false positive rates at a classification threshold of zero. This threshold represents the population mean for each splicing event, providing a biologically interpretable decision boundary that classifies samples based on whether their splicing levels are above or below the cohort average. Cryptic burden was calculated based on the summed z-score cryptic psi across each dataset, including both TDP-43 and non-TDP-43 proteinopathy cases.

## Data and code availability

The RNA sequencing datasets generated in this study are available through ArrayExpress under the accession numbers E-MTAB-15653 and E-MTAB-15433, as well as through the Sequence Read Archive (SRA) under the bioproject PRJNA1256902. The analysis code used in this study is publicly accessible at https://github.com/frattalab/cryptic_biology.git. Code for running the visualization Shiny application is available at https://github.com/frattalab/junction_explorer. The LeafCutter-clustered junction counts for both the NYGC and RiMOD post-mortem datasets can be found at https://zenodo.org/records/15977268. No new resources were generated in this study.

## Contribution

Conceptualization: A-LB, PF

Data Curation: A-LB, MZ, AM

Formal Analysis: A-LB, MZ, AM

Funding Acquisition: PF

Investigation: A-LB, MZ, AM, SB-S, DD, FP, FM, PRM, JNSV

Methodology: A-LB, MZ, AM, PF

Project Administration: PF

Resources: JNSV, JH, MJK, PF

Software: A-LB, MZ, AM

Supervision: A-LB, MJK, PF

Visualization: A-LB, MZ, AM

Writing – Original Draft Preparation: A-LB, MZ, AM, PF

Writing – Review & Editing: A-LB, MZ, AM, SB-S, DD, FP, FM, JH, MJK, PF

## Acknowledgements

A.-L.B. is supported by a Guarantors of Brain Post-doctoral non-clinical fellowship. S.B.-S. was funded by a UK Motor Neurone Disease Association (MNDA) and Masonic Charitable Foundation PhD Studentship (893792) to P.F. This research was funded by a UK Medical Research Council and MNDA Senior Clinical Fellowship and a Lady Edith Wolfson Fellowship (MR/M008606/1 and MR/S006508/1), National Institutes of Health (NIH) U54NS123743 grant and a Target ALS to P.F. M.S. is supported by a UK Research and Innovation Future Leaders Fellowship (MR/T042184/1). P.R.M. is supported by a Wellcome Trust Clinical Training Fellowship (102186/B/13/Z). J.N.S.V. is supported by the Brain Research UK Miriam Marks Fellowship in Neurodegeneration (BF-100029), the Target ALS Springboard Fellowship (FS-2023-SBF-S2) J.H. is supported by NINDS U54NS123743, Target ALS, My Name’5 Doddie, the BrightFocus Foundation, and the Packard Center for ALS Research The funders had no role in study design, data collection and analysis, decision to publish or preparation of the manuscript. The authors would like to thank Maria Secrier for her support and guidance through the initial analyses of these data.

## Conflict of interest

A.M. performs consulting for ISOgenix Ltd. P.F. consults for, holds shares in and is academic founder of Trace Neuroscience. All other authors declare no competing interests.

---

**Extended Data Figure 1.**
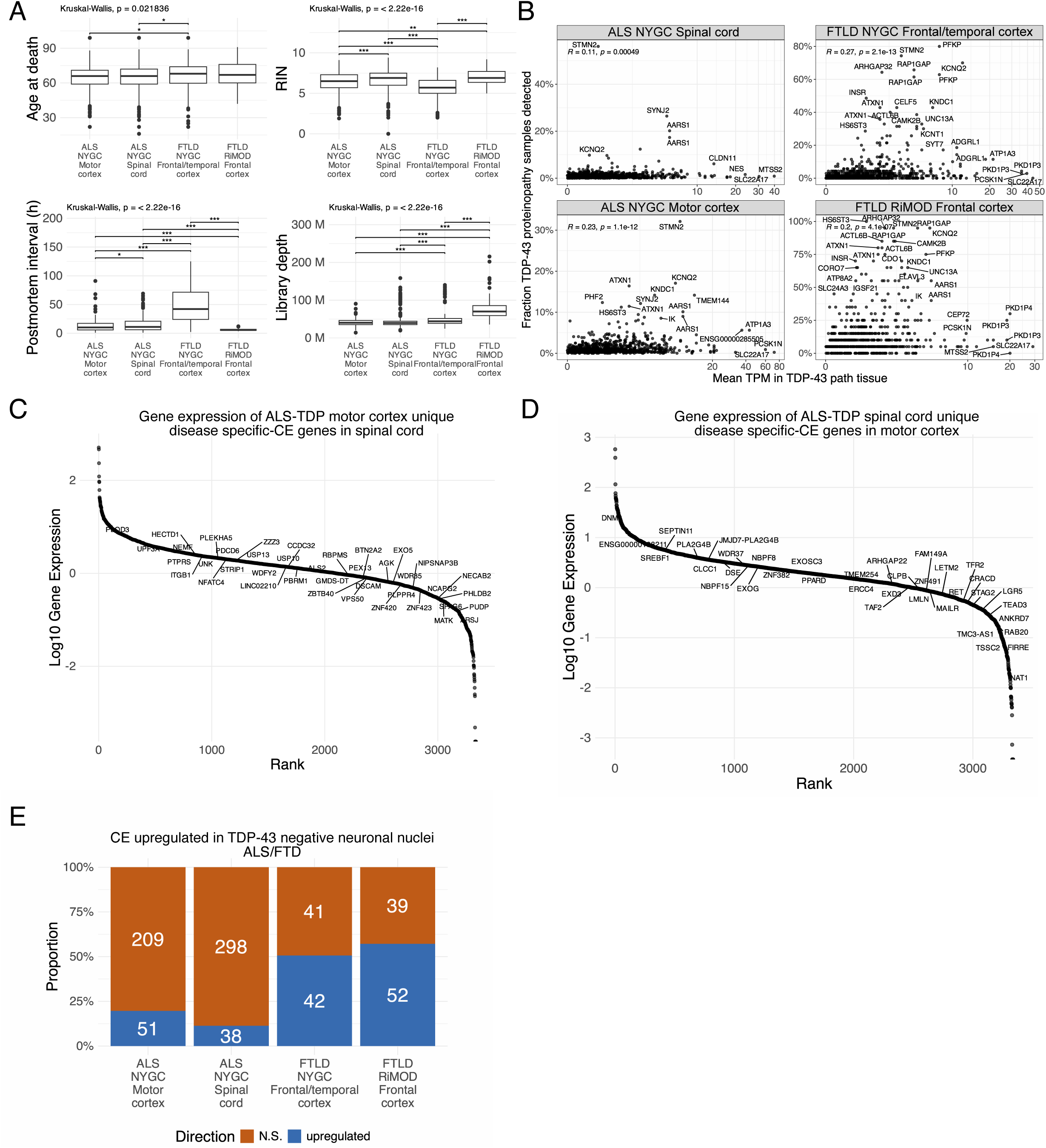
Sequencing features of the postmortem cohorts. A. Age at death (top left), RIN (top right), postmortem interval (bottom left), and library depth (bottom right) in the different cohort analysed (NYGC ALS motor cortex, NYGC ALS spinal cord, NYGC FTLD frontal/temporal cortex, RiMOD FTLD frontal cortex). Wilcoxon test. B. Correlation of the fraction of samples with TDP-43 pathology where a CE is detected (y-axis) and the mean gene expression of that gene in tissues with TDP-43 pathology (x-axis) in the four cohorts (NYGC ALS motor cortex, NYGC ALS spinal cord, NYGC FTLD frontal/temporal cortex, RiMOD FTLD frontal cortex). C. Ranking of genes with motor cortex specific CEs based on their expression in the spinal cord. D. Ranking of genes with spinal cord specific CEs based on their expression in the motor cortex. E. Proportion of TDP-43-proteinopathy specific CE junctions derived from the bulk RNA-seq from NYGC ALS motor cortex, NYGC ALS spinal cord, NYGC FTLD frontal/temporal cortex, and RiMOD FTLD frontal cortex upregulated in TDP-43 negative FACS-sorted neuronal nuclei from FTLD frontal cortex. Box plots: boundaries 25–75th percentiles; midline, median; whiskers, Tukey style. Significance levels reported as * (p<0.05) ** (p< 0.01) *** (p<0.001) **** (p<0.0001).

**Extended Data Figure 2.**
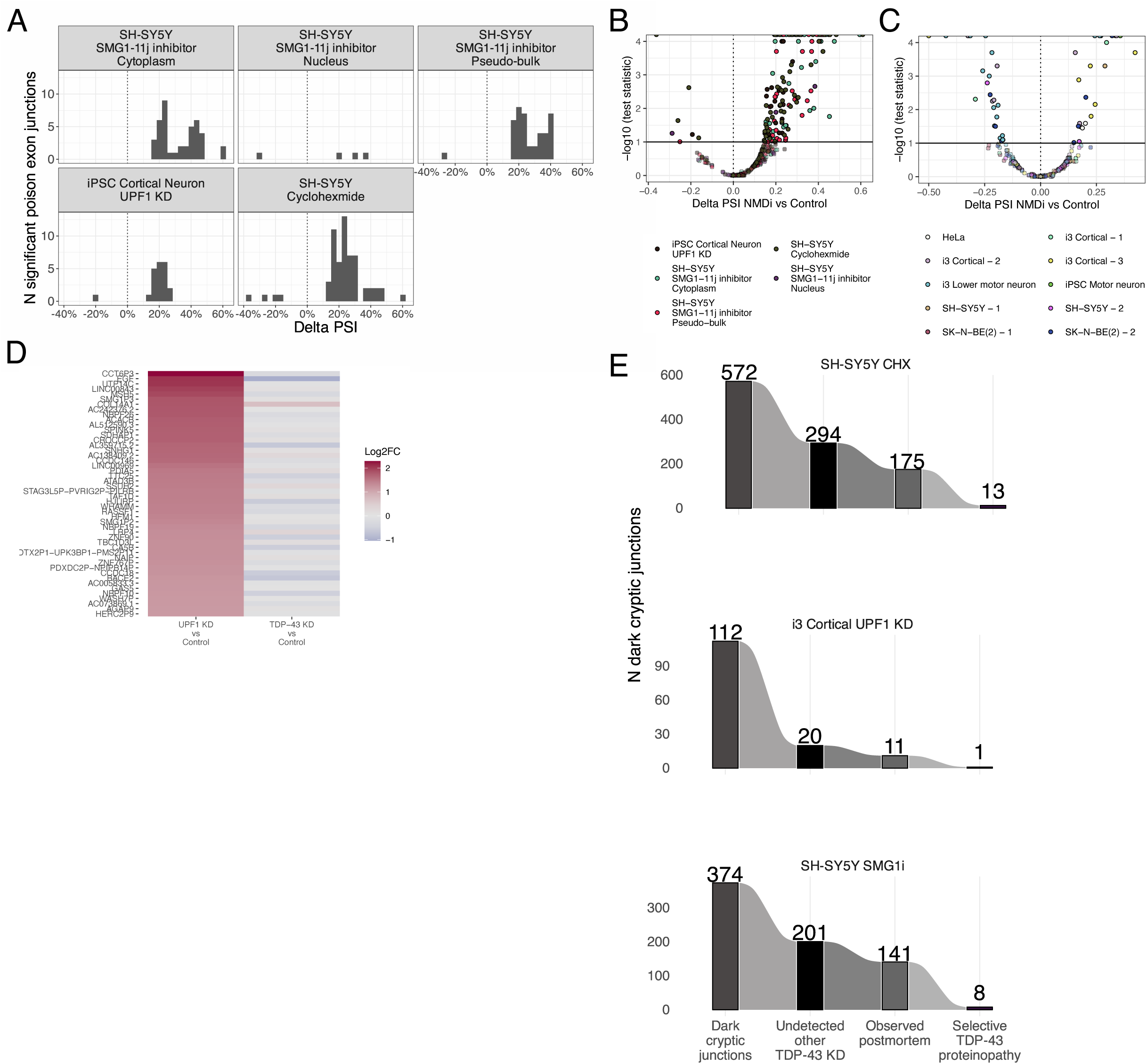
TDP-43 loss does not alter NMD efficiency, but NMD-sensitivity determines CE detection in postmortem tissue. A. Counts of significantly changed poison exon junctions by PSI (3,3% bins) in the different NMD experiments. B. Volcano plot depicting PSI change of poison exon junctions detected in the NMD experiments between NMD inhibition and control condition. C. Volcano plot depicting PSI change of poison exon junctions detected in TDP-43 knockdown experiments. D. Differential gene expression of the 50 most upregulated genes between UPF1 KD and control condition, between UPF1 KD and control condition (left) and between TARDBP KD and control condition (right). E. Number of dark CE, including total number, number undetected in other TDP-43 knockdown, number observed in postmortem tissue, and number selective for TDP-43 pathology, divided by the three NMD inhibition experiments.

**Extended Data Figure 3.**
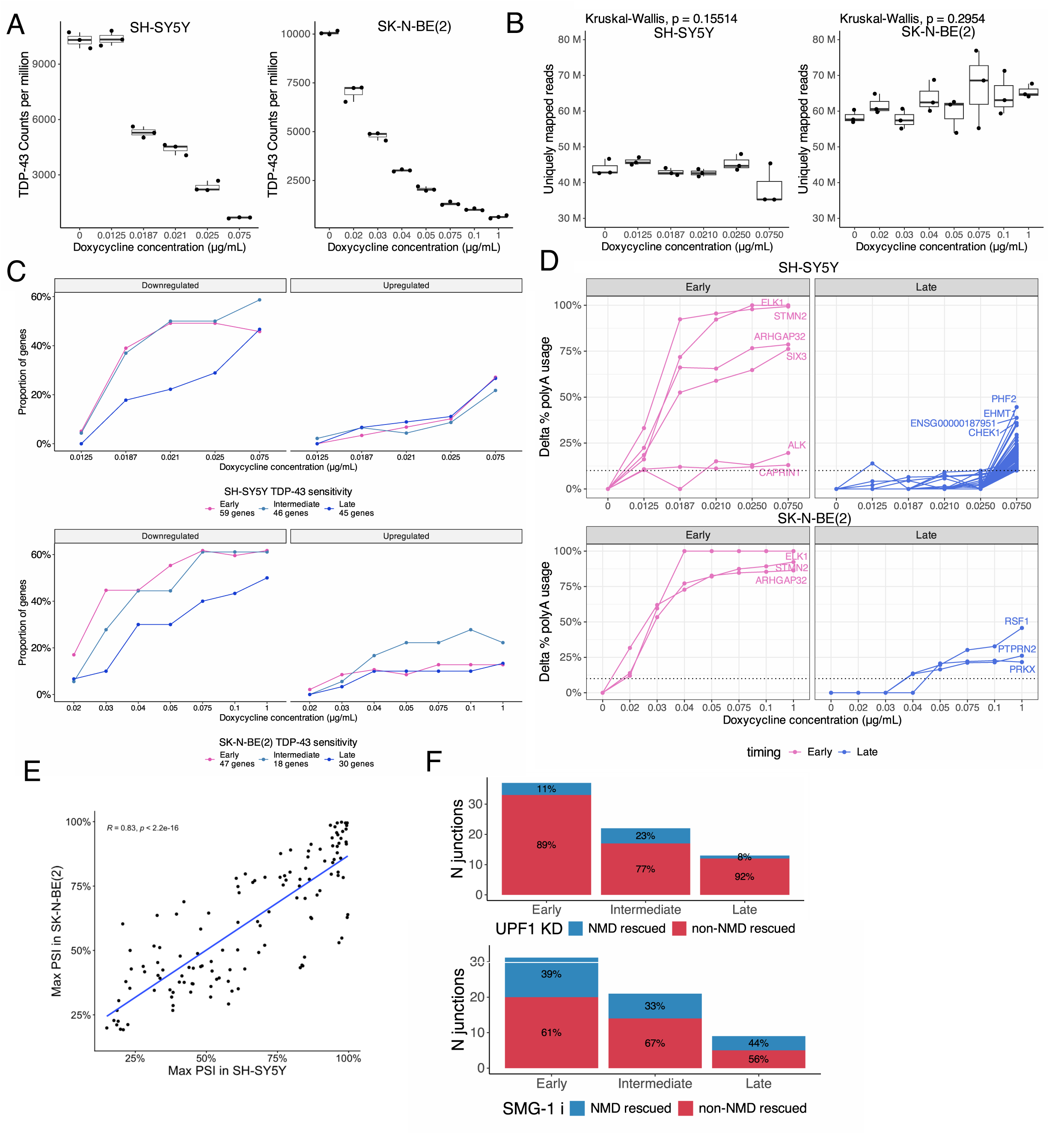
Cryptic splicing sensitivity to TDP-43 levels is reliable across two different neuronal lines and is not driven by sensitivity to NMD. A. TARDBP normalised counts from RNA-seq experiment in SH-SY5Y (left) and SK-N-BE(2) (right) cells. B. Library depth, expressed as uniquely mapped reads, for each doxycycline level in SH-SY5Y (left) and SK-N-BE(2) (right) cells. C. Proportion of genes that are either significantly upregulated (left) or downregulated (right)(adj. p-value < 0.05), split by responsiveness category, in SH-SY5Y (top) and SK-N-BE(2) (bottom) cells. Genes not significant differential not plotted. D. Spaghetti plot showing the response of early (left) and late-responsive (right) last exon cryptic splicing events to TDP-43 loss of function in the SH-SY5Y (top) and SK-N-BE(2) (bottom) cells. E. Scatter plot displaying the PSI at the strongest TDP-43 knockdown level in SH-SY5Y (x-axis) and SK-N-BE(2) cells (y-axis). F. Number of CE junctions detected in each dose-response category split by NMD- (blue) and non-NMD (red) rescue in i3-derived cortical-like neurons with UPF1 KD (top) and SH-SY5Y cells treated with SMGi-j11 (bottom).

**Extended Data Figure 4.**
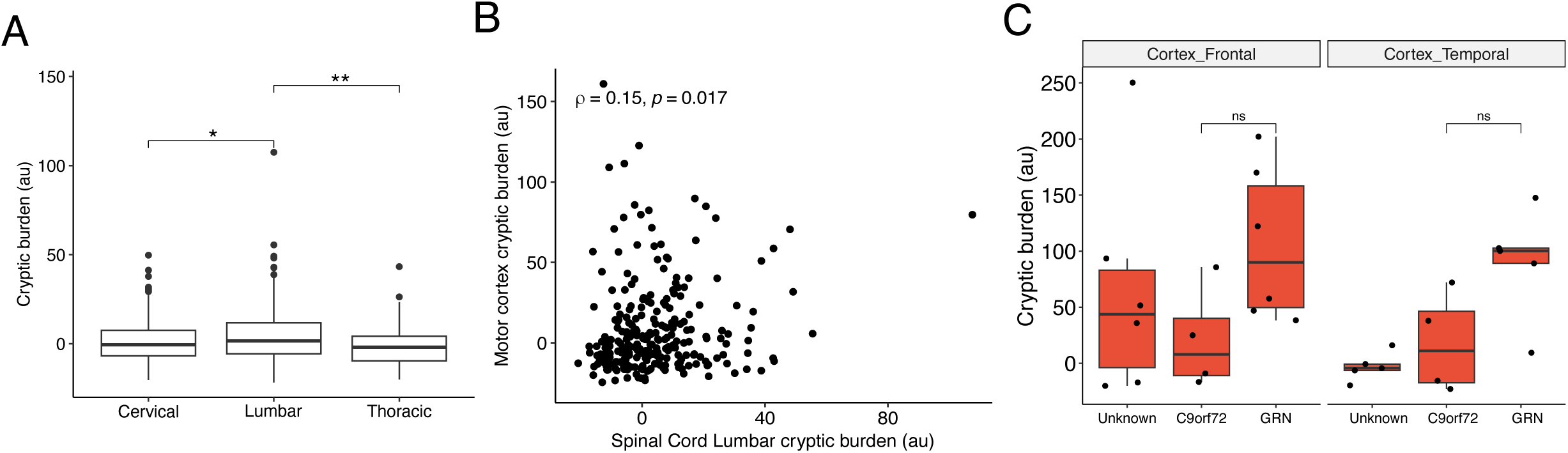
Cryptic splicing burden varies by region and disease mutation. A. Cryptic splicing burden in cervical, thoracic, and lumbar spinal cord from NYCG ALS-TDP samples. B. Correlation between cryptic splicing burden of lumbar spinal cord (x-axis) and motor cortex (y-axis) in the same patients in ALS-TDP. C. Cryptic splicing burden between different genetic types of FTLD (unknown, C9orf72, and GRN) in NYGC FTLD frontal cortex (left) and NYGC FTLD temporal cortex (right), considering only FTLD type A cases.

